# EEG is better left alone

**DOI:** 10.1101/2022.12.03.518987

**Authors:** Arnaud Delorme

## Abstract

Automated preprocessing methods are critically needed to process the large publicly-available EEG databases, but the optimal approach remains unknown because we lack data quality metrics to compare them. Here, we designed a simple yet robust EEG data quality metric assessing the percentage of significant channels between two experimental conditions within a 100 ms post-stimulus time range. Because of volume conduction in EEG, given no noise, most brain-evoked related potentials (ERP) should be visible on every single channel. Using three publicly available collections of EEG data, we showed that, with the exceptions of high-pass filtering and bad channel interpolation, automated data corrections had no effect on or significantly decreased the percentage of significant channels. Referencing and advanced baseline removal methods were significantly detrimental to performance. Rejecting bad data segments or trials could not compensate for the loss in statistical power. Automated Independent Component Analysis rejection of eyes and muscles failed to increase performance reliably. We compared optimized pipelines for preprocessing EEG data maximizing ERP significance using the leading open-source EEG software: EEGLAB, FieldTrip, MNE, and Brainstorm. Only one pipeline performed significantly better than high-pass filtering the data.

## Introduction

Electroencephalography (EEG) is a relatively low-cost brain imaging modality that allows the collection of large quantities of data, and automated preprocessing methods are critically needed to process the large publicly available EEG databases.^1^ Yet, EEG is noisy and often contaminated by artifacts from the environment or the participants. Participants’ eye movements and face, jaw, and neck muscle contractions create scalp electrical potentials about 10 times the amplitude of brain signals and need to be removed.^2^ One strategy to remove noise is the repeated presentation of stimuli in event-related potential paradigms.^3^ A complementary strategy is to use digital signal processing to remove artifacts.^2,4,5^

The currently accepted method recommended by most software for removing EEG artifacts is the visual inspection of the raw EEG data by expert EEG researchers. This procedure is both a time-consuming and imprecise process. There is no consensus on what an EEG artifact is, so one researcher’s data cleaning might differ from another. One reason is that there is large inter-subject variability in EEG data, so the amplitude of the EEG signal can vary widely from one participant to the next. Ideally, multiple researchers would clean the same data, and their aggregated choice would be considered for cleaning the data. This inter-rater agreement is the gold standard for EEG data rejection.^6^

However, inter-rater agreement is difficult to implement. Cleaning EEG data manually on multiple subjects takes several days. To our knowledge, only one dataset with marked rejections from multiple raters has been released.^7^ We used this dataset to determine which automated artifact detection method yielded the best result and found that the chosen algorithm agreed more with each human than humans agreed between themselves.^7^

While waiting for more of these manually labeled collections of datasets to test automated preprocessing and cleaning pipelines, we can use other methods to assess preprocessing data quality, namely, the amplitude of brain response to experimental conditions.^3^ Multiverse analyses scanning the space of preprocessing parameters are a step in that direction, although they still rely on pre-defined ERPs.^8,9^ In this article, we developed a general method to assess the statistical power of detecting a difference in brain response between stimuli. We used this method to determine how filtering, referencing, artifact rejection in different software, and baseline removal influence statistical significance in EEG data. We finally designed and compared a collection of automated pipelines in different open-source software packages.

## I. Results

We assessed the percentage of significant EEG channels between two conditions of interest using randomly resampled trials with replacement with collections of 50 trials per condition (see Methods; see diagram in Supplementary Figure 1) and analyzed data from three publicly available experiments (Supplementary Table 1). The first experiment (Go/No-go) is a go-no visual categorization task where participants are instructed to respond when they see an animal in a briefly flashed photograph. In this task, we compared the brain-evoked potentials between correct animal targets (50%) and correct distractors (50%).^10^ In the second experiment (Face), we compared the brain-evoked potentials of familiar and scrambled faces presented to participants who performed a face symmetry judgment task designed to maintain their attention.^11^ In the third experiment (Oddball), participants had to respond to oddball sounds (70 ms of 1000 Hz) interspaced by frequent standard sounds (70 ms at 500 Hz)^12^ (see Methods).

We high-pass filtered the data at 0.5 Hz, extracted data epochs (from −1 to 2 seconds) with no baseline removal, and calculated at each latency the average number of significant channels using random resamples of 50 epochs (see Methods). We then searched for the 100-ms window with the most significance. We found the latency of maximum effect size in each dataset differed (Figure 1): Go/No-go (350 to 450 ms), Oddball (400 to 500 ms), and Face (250 to 350 ms). This range, custom for each dataset, was used to assess the efficacy of different signal preprocessing methods.

**Figure 1.**
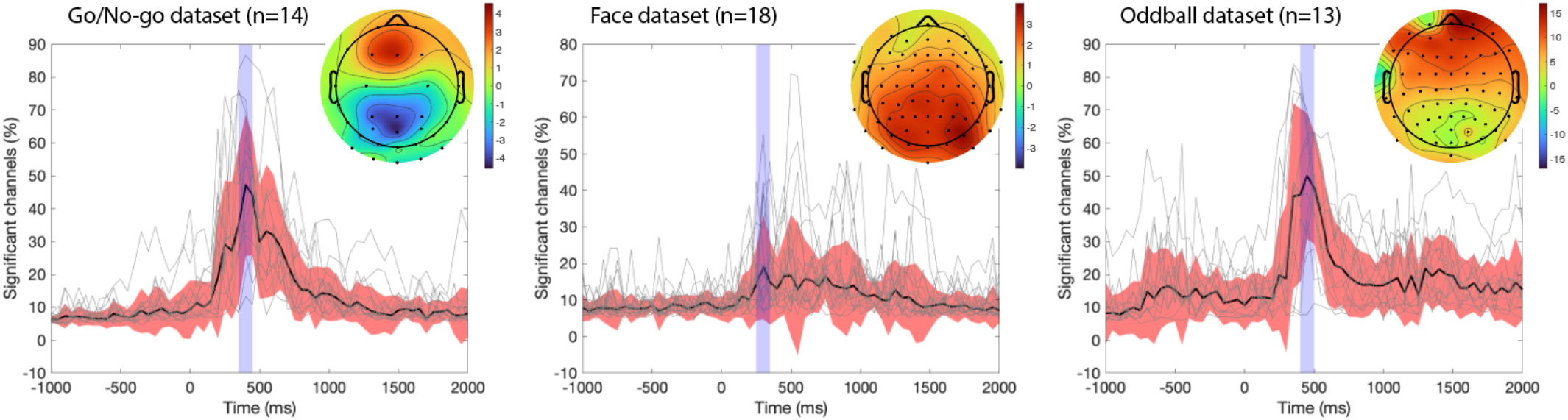
Percentage of significant electrodes across time for three publicly available experiments. Event-related potentials for two conditions (Visual Go/No-go: animal targets vs. non-animal distractors; Face: familiar vs. scrambled faces; Auditory Oddball: oddball vs. standard sounds) are extracted from the raw high-pass filtered data at 0.5 Hz, and the average percentage of significant electrodes in 20,000 random resamples of 50 trials per condition is calculated at each latency in 50-ms increments. Black curves indicate the median, red regions indicate median absolute deviations, and gray curves show the percentage of significant channels for individual subjects in each dataset. The blue region indicates the 100-ms region of maximum effect size, which may correspond to response-related activity for the Go/No-go and Oddball datasets. The scalp topography for each experiment is shown using a μV scale.

### A. Preprocessing methods

#### 1) High-pass filtering

Is there an ideal frequency cutoff to high-pass filter EEG data? Linear filter length increases as the cut-off frequency decreases, making it impractical to filter EEG at frequencies below 0.5 Hz. As a result, we used a 4th-order Butterworth filter to assess the optimal high-pass filter cutoff frequency (ERPLAB package; see Methods). We tested filters at 0.01 Hz, 0.1 Hz, 0.25 Hz, 0.5 Hz, 0.75 Hz, and 1 Hz. Filtering had the most important effect on the percentage of significant channels compared to all other preprocessing steps. It improved performance by 13% (Face), 47% (Oddball), and 57% (Go/No-go). The best filtering was at 0.1 Hz (Face), 0.5 Hz (Oddball), and 0.75 Hz (Go/No-go). Filters above 0.1 Hz led to significant improvement compared to no filtering for the Visual Go/No-go and Auditory Oddball datasets (p<0.0001), and also led to significant improvement compared to the 0.01-Hz filter (p<0.0001). Although there was a significant difference between a filter at 0.01 Hz and one at 0.5 Hz for two datasets (Go/No-go and Oddball; p<0.0001 in both cases), this was not the case for the Face dataset, likely because of a filter applied during EEG acquisition (see Discussion).

Not all filters are created equal. Most software packages have various options and parameters to design filters, and it was impractical to test them all. We tested the default filter in the publicly available software packages EEGLAB, MNE, Brainstorm, and FieldTrip (see Methods and Supplementary Figure 2) and compared them to the reference filter in Supplementary Figure 2. There ERPLAB reference filter performed better than all other filters for the Oddball dataset (p<0.0001) but not for the Face and Go/No-go dataset. For the Go/No-go dataset, the MNE filter performed significantly worse than the ERPLAB reference filter (p<0.0001), the Brainstorm filter (p<0.005), and the EEGLAB filter (trend at p=0.02). The MNE filter performed worse for the Face dataset than the EEGLAB filter (p<0.002). For Fieldtrip, we tried two methods: one-step preprocessing, which we realized filter raw data epochs, and multi-step preprocessing, which allows filtering raw data before extracting data epochs. For the Oddball dataset, the FieldTrip filter applied to data epochs decreased the percentage of significant electrodes up to 25% compared to the reference filter (p<0.0001) and should be avoided. We use the default EEGLAB filter (see Methods) to high-pass the data from all datasets at 0.5 Hz in all analyses in section A.2 to A.5 -- including those using the MNE, Brainstorm, and FieldTrip software packages.

#### 2) Line noise removal

There are several ways to remove or minimize line noise in EEG data (see Methods; Supplementary Figure 3). The most common method is filtering out the line noise frequency using a band-stop filter, also known as a notch filter. We used an FIR notch filter and IIR notch filter (see Methods), and observed no change in the percentage of significant channels for any of the datasets - note that the Visual Go/No-go dataset already had the line noise notched (Supplementary Table 1). We also tested the *cleanline* EEGLAB plugin^13^, which estimates and removes sinusoidal line noise, and the *Zapline-plus* EEGLAB plugin^14^, which combines spectral and spatial filtering to remove line noise. These methods did not yield significant differences compared to no line-noise removal, except for the Face dataset where they led to a small yet significant performance decrease (*cleanline* p<0.001) or a trend performance decrease (*Zapline-plus* p<0.02). We also rejected noisy channels based on their activity distribution at the line noise frequency (see Methods). We interpolated channels that had 1, 1.25, 1.5, 2, 3, and 4 standard deviations more line noise than other channels (see Methods). For the Go/No-go dataset, rejecting noisy channels led to significant performance improvements (p<0.002 for all standard deviations values tested), with a 1 standard deviation threshold leading to the best results for the Face dataset (average of 17% of channels rejected) and a 1.25 standard deviation threshold for the Auditory Oddball dataset (average of 17% of channels rejected). For our final pipeline (see section B), we chose the default threshold of 4 standard deviations, which led to significant improvement for the Face and Oddball datasets (p<0.005 in both cases) and a modest percentage of channels rejected (average of 1% to 6% depending on the dataset).

#### 3) Referencing

Figure 3 shows the comparison of using different references for the Oddball dataset, for which it was possible to test more reference montages than for other datasets (see Supplementary Figure 4 for the Go/No-go and Face data). For this dataset, all re-referencing significantly decreased the percentage of significant channels (p<0.005 in all cases), including the PREP reference, which uses channel interpolation to remove artifacts (see Methods). For the Go/No-go dataset, a trend decrease in performance was observed for all reference types (p<0.09 or lower), and a non-significant decrease was also noted for the Face dataset. For the Oddball dataset, the median, average, REST, and PREP references decreased performance the least, although they were not significantly better than other reference methods except the circumferential reference. The Nose reference introduced the highest variability (Figure 3). The circumferential reference daisy chain montage also performed significantly below the median, average, REST, and PREP references for the Go/No-go, Face, and Oddball datasets (significance or trend of p<0.02 in all cases).

**Figure 2.**
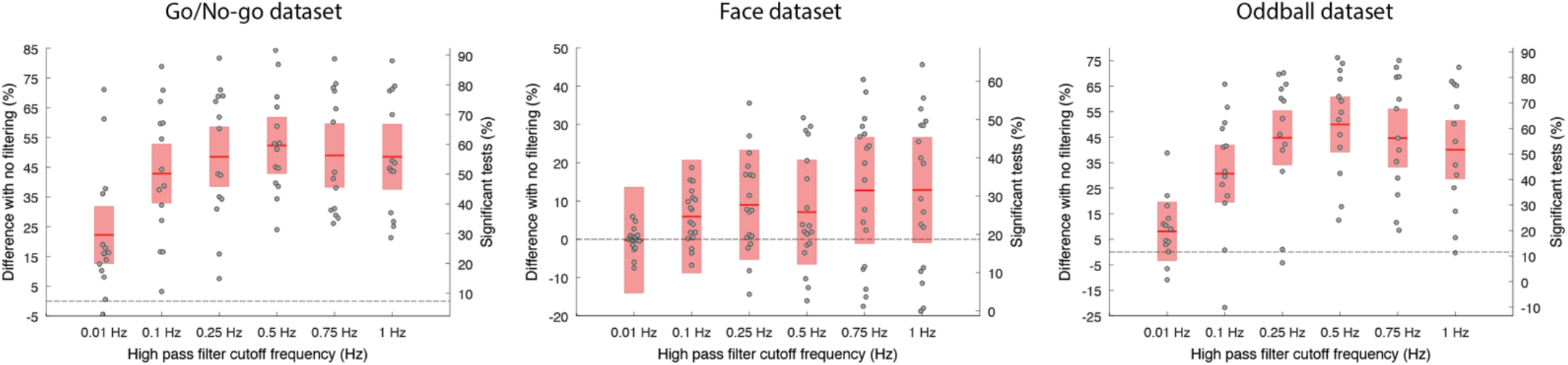
Influence of high-pass filtering cutoff frequency on the percentage of significant channels compared to no filtering for three datasets. The red regions indicate the 95% confidence interval, and the dots indicate individual subjects. The ordinate scale on the right of each panel indicates the percentage of significant tests. It is obtained by shifting the left ordinate scale by the average percentage of significant channels in the nofiltering condition.

**Figure 3.**
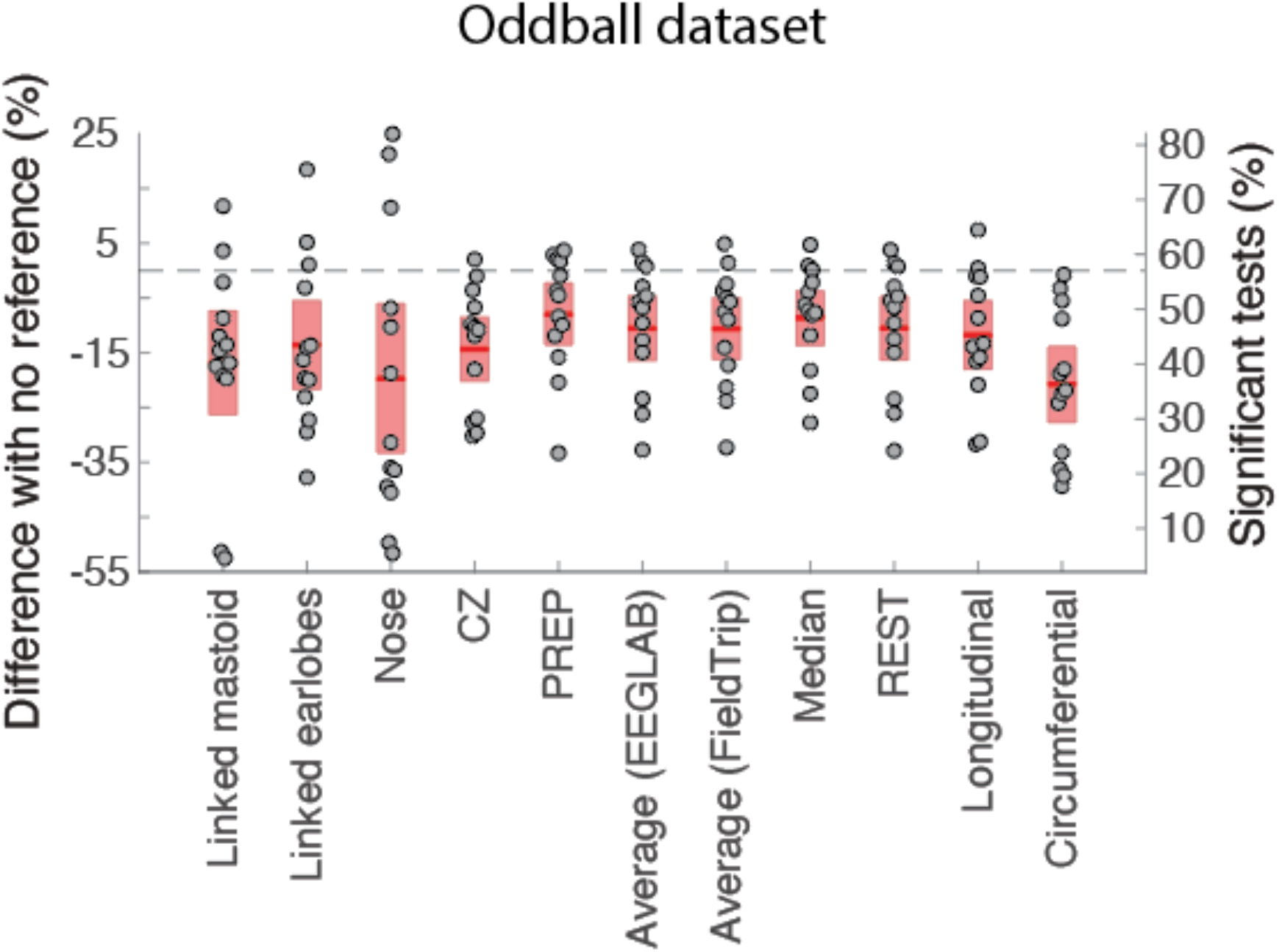
Influence of the reference on the number of significant channels compared to internal BIOSEMI reference (CMS/DRL). For all references, the raw data is first high-pass filtered at 0.5 Hz (see Methods). The red regions indicate the 95% confidence interval, and the dots indicate subjects. See Figure 2 for details regarding the ordinate scales.

#### 4) Artifact rejection methods

We used the same method as in sections A.1-A.4 to assess the performance of automatic EEG artifact rejection which did not remove data trials, such as EEGLAB *clean_rawdata* channel rejection and the EEGLAB *ICLabel* eye movement and muscle rejection. However, for methods removing trials, we had to compute significance using all trials instead of resampling 50 trials, since our goal was to assess whether these methods can increase the percentage of significant channels after bad trials are removed (see Methods).

##### EEGLAB clean_rawdata channel rejection

The core plugin *clean_rawdata* was used for detecting bad channels with correlation thresholds ranging from 0.15 to 0.975 (see Methods and Supplementary Figure 5A). A higher correlation threshold is more aggressive at rejecting channels. We found that a correlation of threshold of 0.95 led to a significant 2% improvement compared to no channel rejection for the Face dataset, with 7% of channels rejected. The best correlation was 0.97 for the Oddball dataset, with 29% of channels rejected and a 15% increase in performance. To avoid rejecting too many channels, for the final EEGLAB pipeline (section B), we chose to use 90%, which led to a significant performance increase for the Face and Oddball datasets (p=0.003 and p<0.0001 respectively) and rejected between 3% and 12% of the channels, depending on the dataset.

##### EEGLAB clean_rawdata ASR rejection

The core plugin *clean_rawdata* was used for detecting bad data segments using the first step of the Artifact Subspace Reconstruction (ASR) method^15^ (see Methods and Supplementary Figure 5B). The threshold used for ASR ranged from 5 to 200 (Table 1). No ASR value led to a significant increase in performance. A low threshold of 5 tended to reject 100% of the trials for all subjects n the Oddball dataset. Since ASR rejection did not significantly decrease the percentage of significant channels, for the final EEGLAB pipeline (section B), we chose the default value, which is equal to 20.

**Table 1.**
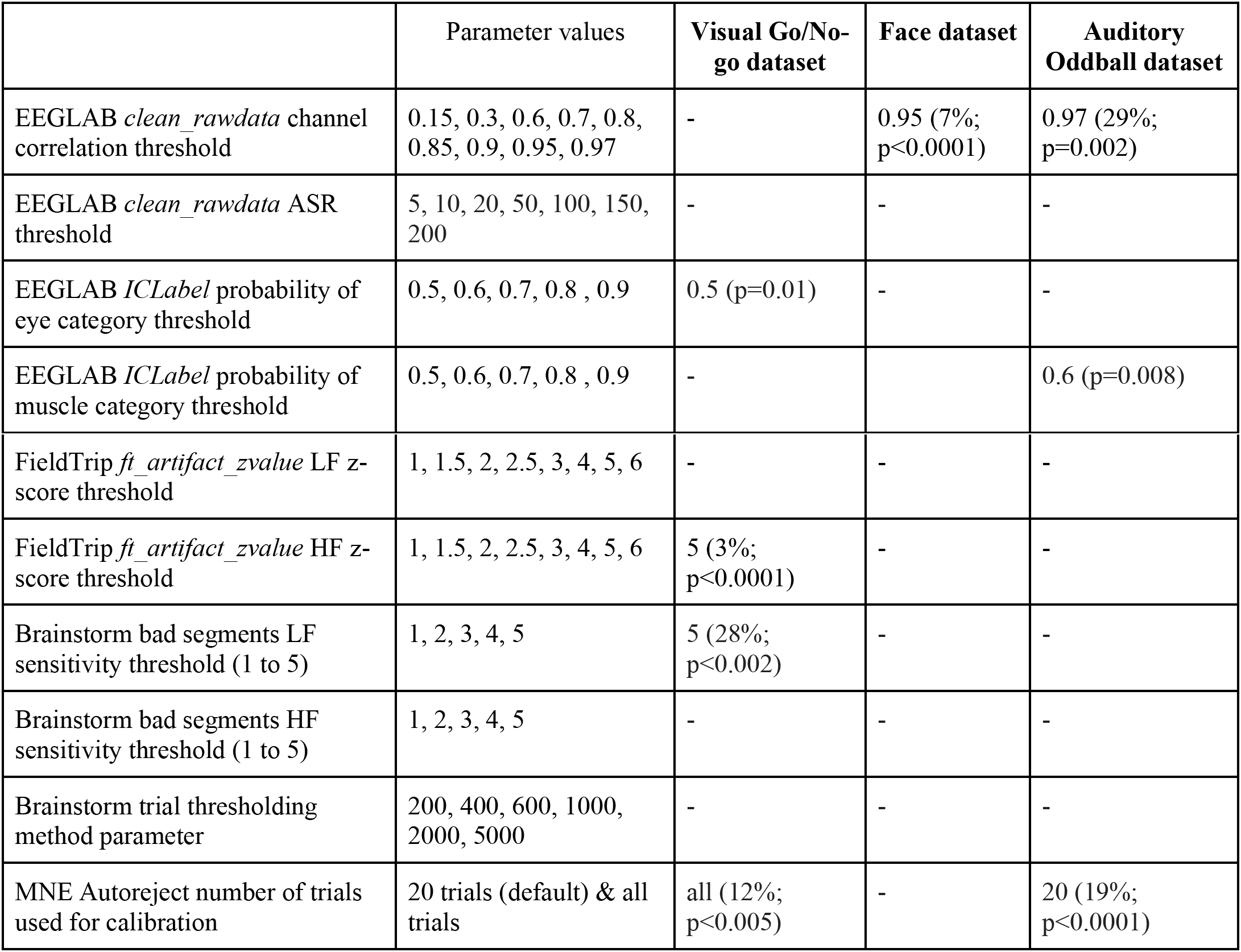
Evaluation of different methods for automated artifact rejection in the most popular open-source software packages for EEG data analysis (EEGLAB, FieldTrip, Brainstorm, and MNE). For all data rejection methods, the data is first high-pass filtered at 0.5 Hz (see Methods). The percentage of significant channels is compared to raw data high-pass filtered at 0.5 Hz. For each method, we indicate the range of parameters tested. Then for each dataset, the first value indicates the optimal parameter, the second value (in parentheses) is the percentage of rejected trials (or interpolated channels for the EEGLAB clean_rawdata channel correlation method) followed by the p-value (see Methods).

##### EEGLAB ICLabel eye movement and muscle rejection

Independent Component Analysis (ICA) was first applied to EEG (*Picard* plugin; see Methods and Supplementary Figure 6). *ICLabel*^16^ was then applied to detect artifacts with thresholds ranging from a probability of 0.5 to 0.9 for belonging to the eye or muscle component category. For eye artifacts, we found a trend advantage of using ICA for the Go/No-go dataset (p<0.05 for all thresholds from 50% to 90%, with a minimum of p=0.01 at 50%). For muscle artifacts, we found a significant advantage to using ICA for the Oddball dataset (0.6 threshold and p=0.008). For the final EEGLAB pipeline (Section B), we used the default value, which is 0.9. We also tested *ICLabel* on other ICA algorithms commonly used on EEG data^17^ (*runica, AMICA, FastICA*, and *SOBI*), but none increased performance significantly (Supplementary Figure 7).

##### FieldTrip ft_artifact_zvalue

The function *ft_artifact_zvalue* of FieldTrip was used to detect artifacts automatically (see Methods). We varied the z-score threshold from 1 to 6 (Table 1 and Supplementary Figure 8). For low frequency, we observed no significant performance improvement. A threshold below 4 for the Oddball dataset decreased performance significantly (p<0.0005). For high frequency, a threshold of 5 or 6 increased performance for the Go/No-go dataset (p<0.0001) but decreased performance for the Oddball dataset (p<0.001). In the final FieldTrip pipeline (Section B), we set the threshold to 4 for both low and high frequency, as recommended in the FieldTrip tutorial (see Methods).

##### Brainstorm bad segment detection

In Brainstorm, we used the function to detect bad data portions (see Methods and Supplementary Figure 9). This function allows finding low-frequency or high-frequency artifacts with a sensitivity level varying from 1 to 5. For low frequency, a sensitivity of 5 led to a significant increase in performance for the Go/No-go dataset (p<0.002). Sensitivity values of 1 and 2 also led to a significant performance increase (p<0.003) but rejected one or more subjects. There were no significant differences for other datasets, except for the sensitivity value of 1 in the Oddball dataset, which rejected 11/14 subjects. Given the aggressiveness of this method, we used the lowest sensitivity of 5 in the final Brainstorm pipeline. For high-frequency artifacts, at levels 1 and 2, no trials were left for any subjects in any datasets. Level 5, the least aggressive sensitivity value, provided trend improvements for the Oddball dataset (p<0.005), with 19% of trials rejected. We used this threshold in the final Brainstorm pipeline (section B).

##### Brainstorm bad trial detection

In Brainstorm, we also used the function to detect bad data trials (*process_detectbad*), varying the threshold from 200 to 5000 (see Methods and Supplementary Figure 9). No value led to a significant increase or decrease in performance. We used a threshold of 200 in the final Brainstorm pipeline (section B).

##### MNE Autoreject

We used MNE and the *Autoreject* plugin library (see Methods and Supplementary Figure 9). *Autoreject* scans the parameter space for optimal values and the main free parameter is the number of epochs to fit the data. We used 20 epochs and all epochs (from about 300 to 600 epochs depending on the dataset) (Supplementary Figure 10) and observe no significant difference between the two. In the optimal MNE pipeline (section B), we fitted the data using the first 20 epochs to speed up computation, as advised in the *Autoreject* tutorial (see Methods).

#### 5) Baseline

We tested if pre-stimulus baseline periods ranging from 100 ms to 1000 ms affected the percentage of significant channels (Figure 4). Applying a pre-stimulus baseline did not significantly improve performance for any of the three datasets. In fact, all baseline tests performance significantly decreased or trended - p<0.025 in all cases except for baseline ranging from 500 ms to 1000 ms in the Oddball dataset. Performance decreased the most for shorter baselines, with a significant performance drop of 3% to 6% for the 400-ms baseline in all datasets (Figure 4) and a significant performance decrease between the 400-ms and the 1000-ms baseline for the Go/No-go and Oddball datasets (p=0.001 in both cases).

**Figure 4.**
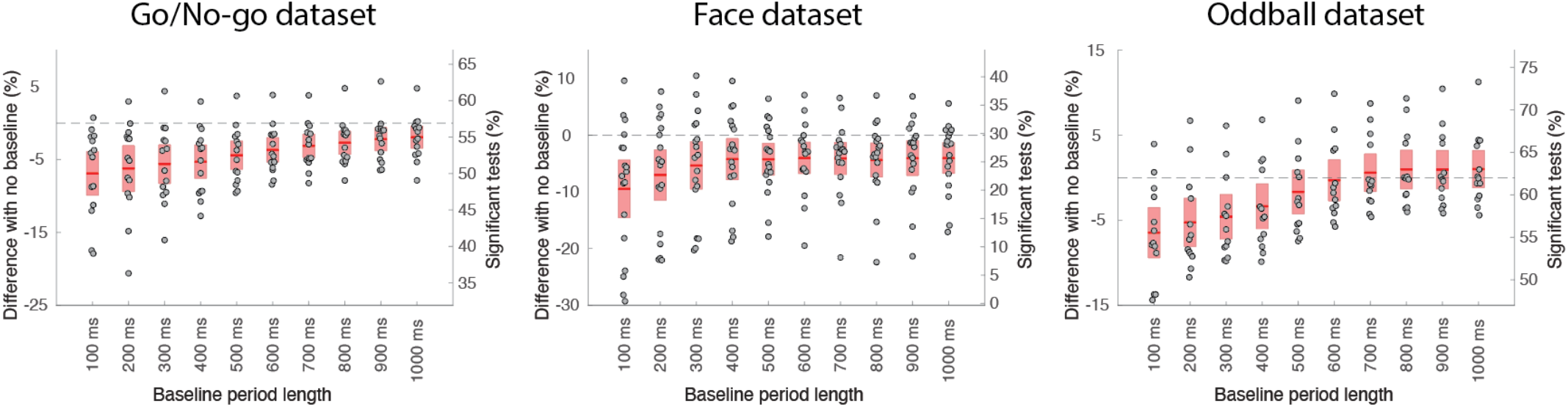
Influence of the baseline period on the number of significant channels compared to no baseline. The red regions indicate the 95% confidence interval, and the dots indicate individual subjects. See Figure 2 for details regarding the ordinate scales.

These results contradict standard practice in event-related potential (ERP) research, although the results might have been different if the data had not been high-pass filtered at 0.5 Hz. To address this issue, we compared two approaches. We compared using a filter at 0.01 Hz with a 200-ms pre-baseline interval (baseline method), an approach proposed by Luck and collaborators,^18^ with a filter at 0.5 Hz and no baseline (Supplementary Figure 10). For all three datasets, using the baseline method led to a significant and large decrease in performance of 30% for the Go/No-go dataset (p<0.0001), 8% for the Face dataset (trend at p=0.02), 42% for the Oddball dataset (p<0.0001) (Supplementary Figure 11). As indicated in Supplementary Table 1, the Face dataset was high-pass filtered at 0.1 Hz at recording time, and the decrease of 8% in performance we observed would likely be higher if this had not been the case.

### B. Optimized pipelines

For each open-source software package, we built an optimal pipeline based on results presented in A.4. Section A.4 indicates how we chose parameters for each artifact rejection method. At the end of the preprocessing pipelines, epochs are extracted, and performance is computed (see Methods). Unlike sections A.1-A.4 for which EEGLAB was always used to filter the raw data and extract data epochs, all pipelines were implemented stand-alone in each software package, meaning that code from other toolboxes was not used. The goal is to provide an optimal integrated and self-contained pipeline to the users of each software package – pipelines are available publicly on GitHub (https://github.com/sccn/eeg_pipelines). These pipelines are optimized to maximize the percentage of significant channels, so no referencing (section A.3) and no baseline (section A.5) are applied (see Methods).

Despite using the most conservative available settings to reject artifacts, the Brainstorm pipeline rejected all trials for some subjects (Figure 5; Go/No-go: n=4; Face: n=14; Oddball n=1). This is likely because Brainstorm is tailored for MEG processing, where artifacts’ amplitude might differ. The FieldTrip pipeline failed to process some datasets (Go/No-go: n=3; Face: n=4) for which time-locking event latencies were not falling on data samples. Both the FieldTrip and Brainstorm tool developers have been notified of these issues, and we expect they will fix them.

**Figure 5.**
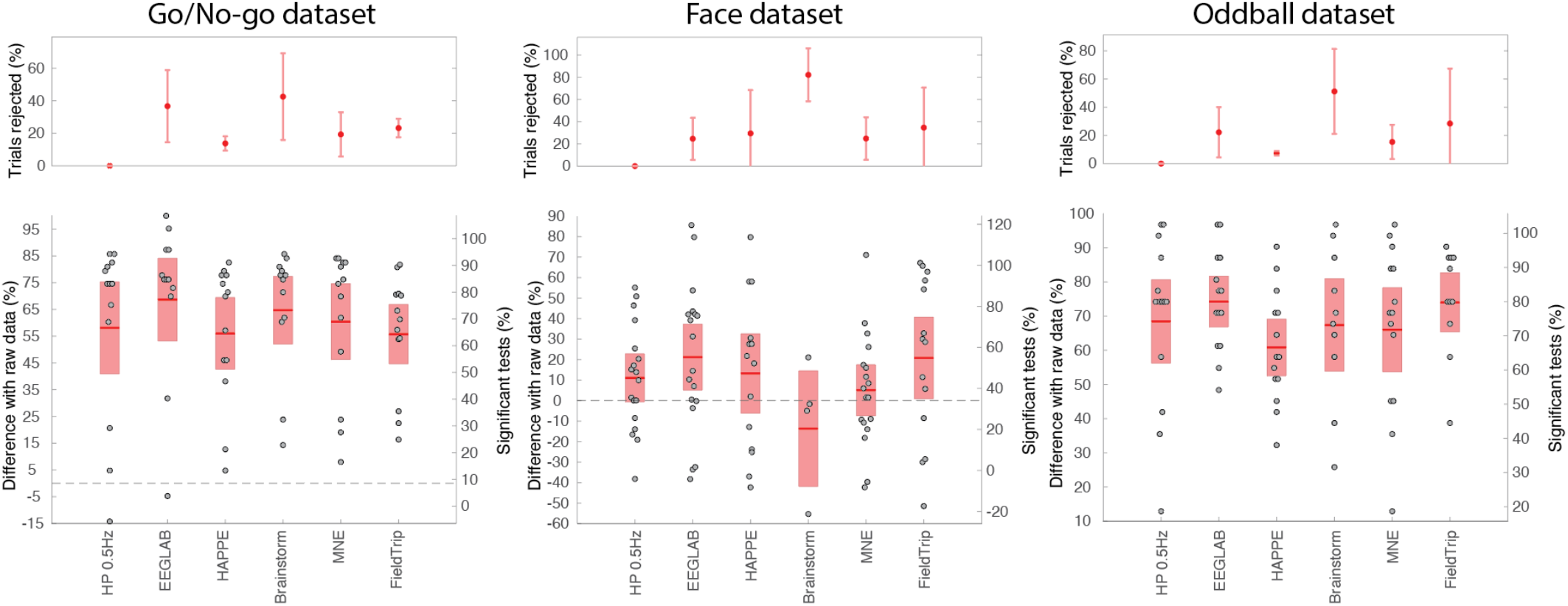
Different automated pipeline percentage of significant channels compared to unprocessed raw data for three datasets. The upper panels show the percentage of trials rejected and the standard deviation. The lower panels show the percentage of significant channels compared to unprocessed raw data. The red regions indicate the 95% confidence interval, and the dots indicate individual subjects. See Figure 2 for details regarding the ordinate scales.

For the Go/No-go and Oddball datasets, all pipelines led to a higher percentage of significant electrodes than no preprocessing (p<0.0001). The EEGLAB pipeline performed best overall. It performed better than the high-pass 0.5 Hz pipeline for the Face (p<0.004) and the Oddball dataset (p=0.0008). The HAPPE, Brainstorm, Fieldtrip, and MNE pipelines did not perform significantly better than the high-pass 0.5 Hz pipeline for any of the datasets. The EEGLAB pipeline performed significantly better in pairwise comparisons with all other pipelines for the Face dataset (p<0.006 except FieldTrip), and it performed better than all other pipelines for the Oddball dataset (p<0.005 except Brainstorm). Fieldtrip performed significantly better than HAPPE (p=0.01) for the Go/No-go dataset. We noted no other pairwise significant comparison.

We did not implement fine-tuning for any pipeline. It is important to note that we used the recommended values for most pipelines (see Methods). Exceptions were as follows. (1) For EEGLAB, channel correlation was set to 0.9 instead of the 0.85 default (Section A.4). (2) For HAPPE, we used a 150-microvolt threshold. (3) For Brainstorm, the sensitivity was decreased to 5 instead of 3 (default) for both low and high-frequency artifact rejection, and trial rejection was set to more conservative −200 to 200 instead of - 100 to 100 (default); For FieldTrip and MNE, we use all default parameter values (see Methods).

## II. Discussion

In a previous report, we used the percentage of data rejected by *clean_rawdata*^15^ and the number of brain components isolated by *ICLabel*^4^ as indicators of data quality.^1^ However, since they are part of existing processing pipelines, these methods cannot be used as ground truth to assess the true quality of EEG data and benchmark pipelines.

In this report, we used a data metric inspired by previous work,^3^ where researchers counted the number of significant trials necessary to reach significance in an oddball paradigm. However, this was performed on a single electrode, and we observed that results would vary widely based on the reference chosen. Because of volume conduction, a strong differential effect between conditions may be visible and significant on most electrodes, so we chose to compute significance on all electrodes. Figure 1 confirms this hypothesis: for several subjects, more than 80% of their electrodes showed significance in the window of interest. Our procedure is also agnostic to the choice of reference and does not depend on the number of trials or channels (see Methods). Instead of looking at differences between conditions, it would also have been possible to test for absolute ERP amplitude differences compared to baseline. However, maximizing the ERP amplitude and the significance on multiple channels might not be cognitively relevant. By comparing conditions, our quality metric is guaranteed to be relevant to researchers.

Filtering increased the percentage of significant channels by about 50% for the Go/No-go and Oddball datasets. For the Face dataset, filtering led to non-significant improvement, likely because of the high-pass filter at 0.1 Hz at the time of data collection (Supplementary Table 1). When comparing software implementations, the performance was originally low for the FieldTrip filter in the Oddball dataset (Supplementary Figure 2). We realized that the FieldTrip preprocessing function extract data epochs before filtering the data when provided with both filter settings and epoch information. After consulting with FieldTrip developers, for all analyses using FieldTrip, we used an alternate multi-step implementation allowing to extract epochs after the raw data had been filtered.

We found that the only line noise removal that affected the percentage of significant channels was the interpolation of noisy channels. Notch filtering had no significant effect. Spectral and spatial interpolation in *cleanline* and *Zapline-plus* even impacted performance negatively for the Face dataset. The Go/No-go data likely had a hardware notch filter that proved efficient at removing line noise since removing noisy channels no longer affected performance (Supplementary Table 1; Supplementary Figure 3). When significant, the performance improvement of line noise correction was minor (on the order of a few percent), in contrast to the effect of high-pass filtering, which led to performance increases of about 50% on two datasets. Except for data visualization and cluster corrections for multiple comparisons in the frequency domain, offline removal of line noise might not be a critical step when preprocessing EEG data.

We have found that re-referencing in all three datasets did not increase the percentage of significant channels. At best, the median, average, REST, and PREP references led to a non-significant decrease in performance for some datasets. Again, this is contrary to belief in the EEG community, where there is a relentless search for the optimal EEG reference.^19–21^ Our result is surprising, given that the three datasets used different references (Cz for the Go/No-go data, nose for the Face data, and BIOSEMI internal reference for the Oddball data). The nose reference for the Face dataset has been known to introduce artifacts, especially at high frequency,^22^ and is now seldom used in EEG research. The BIOSEMI raw signal for the Oddball dataset is considered unsuited for offline data analysis, and the BIOSEMI company recommends choosing an offline reference to add 40 dB extra CMRR (common mode rejection ratio) (https://www.biosemi.com/faq/cms&drl.htm). The Neuroscan Cz reference for the Go/No-go dataset is also a common reference by EGI, Inc. Nevertheless, re-referencing the data to Cz for the Oddball dataset did not increase performance (Figure 3). These results might indicate a fundamental difference in terms of noise suppression between analog referencing and digital re-referencing and warrant more research to understand the difference between the two. The interaction between data re-referencing and prior data filtering (0.5 Hz high-pass in our case) should also be investigated.

We have observed that, if the data is high-pass filtered at or above 0.5 Hz, subtracting mean baseline activity should be omitted for event-related analyses. Subtracting baseline activity has either no effect or a negative effect on data quality, especially when the baseline is shorter than 500 ms. Lutz and collaborators argue that filtering distorts ERPs and that baseline removal should be used instead.^18^ While it is true that the ERPs are distorted, we have found that, when using the short baseline proposed by Lutz, data quality decreases dramatically, rendering almost all electrodes non-significant for some subjects in the case of the Oddball dataset (Supplementary Figure 10).

Regarding artifact rejection, our results are not what one would expect of basic (simple thresholding) and more advanced data cleaning methods. Almost none of them led to significant performance improvement, and when improvement was observed, the effect was weak and not consistent across datasets. When resampling 50 data trials instead of using all the trials to compute significance, we observed that rejecting trials led to significant improvement for most methods, meaning that these methods are indeed capable of removing bad trials. For example, using a 50 data trials bootstrap with ASR, a threshold of 10 increased performance for the Face and Oddball datasets (p<0.0001 in both cases), although about 60% to 80% of trials were rejected. Using a 50 data trials bootstrap, Brainstorm sensitivity 5 for high-frequency artifacts provided significant improvements (p<0.004 in all datasets with 19% to 45% of trials rejected), and MNE Autoreject also provided significant improvements (Go/No-go: p=0.005 and 12% of trials rejected; Oddball: p<0.001 and 19% of trials rejected). However, as seen in Table 1, this was not the case when bootstrapping all remating trials: the removal of bad trials most often failed to compensate for the decrease in the number of trials and associated decrease in statistical power compared to the control condition where no trials were removed.

Although ICA increased the percentage of significant channels, it did not do so systematically (Table 1). However, there is more to consider when using an artifact rejection method than the number of significant channels. For example, ICA and ICLabel removed artifacts related to eye and muscle movements, which affect scalp topographies and might provide advantages for visualization (Supplementary Figure 12) and subsequent source localization.

The EEGLAB pipeline performed best and was a significant improvement compared to filtering the data for all datasets, with 5% (Go/No-go) to 17% (Oddball) and 18% increase in the number of significant channels. This was likely due to the methods used to reject and interpolate bad channels, as both the ICA and the ASR method showed limited efficacy (section A.4). All other pipelines showed no improvements compared to filtering the data. One could argue that we have a double-dipping problem because we optimized the pipelines, then recalculated significance. To address this issue, we reverted the EEGLAB pipeline^1^ to all default parameters (v2022.0), which only changed the threshold for channel correlation to 0.85 instead of 0.9. The difference with the basic pipeline was no longer significant for the Go/No-go dataset. It was still significant for the Face (p=0.004) and the Oddball dataset (p=0.0008).

We could not include all existing pipelines in our comparison. The PREP pipeline^23^ is an EEGLAB plugin for building EEG pipelines. The PREP pipeline contains threshold-based automated EEGLAB artifact rejection,^2^ which we chose not to test since we tested similar methods in other software (Brainstorm, FieldTrip). The PREP pipeline also contains a method to perform robust referencing, which we included when comparing re-referencing methods (Figure 3). The APICE pipeline is another EEGLAB pipeline we did not test since it is tailored to infant EEG.^24^ EEGLAB also contains 29 plugins with methods to reject artifacts, including FASTER^25^ and MARA,^26^ two popular automated EEG processing methods. Although we did not evaluate these algorithms directly, they were included in the HAPPE pipeline.

We show the efficacy of using a non-subjective task-relevant metric to assess data quality that may be used to guide EEG preprocessing. Using this metric, we also showed that for relatively clean EEG acquired in laboratory conditions, preprocessing techniques had little effect on data quality. This might not be the case for more noisy data acquired in other conditions.^7^ We hope future work continues to develop the proposed method to unveil a new era of automated signal processing for EEG, MEG, and iEEG.

## III. methods

### Data quality metric

For Figure 1, we high-pass filtered the data at 0.5 Hz (we used the default EEGLAB hamming windowed sinc FIR filter of the Firfilt plugin (v2.6) with 0.5-Hz high-frequency cutoff and default parameters) and extracted 3-second epochs (from −1 to 2 seconds) with no baseline removal. We calculated the number of significant channels in a random resamples of 50 epochs with replacement and with 20,000 repetitions at each latency (in 50-ms increments) for each subject (Supplementary Figure 1). We then calculated the median and median absolute deviation (MAD) at each latency. To find the window of most significance, we smoothed the median values (moving average filter of size 3) and took the maximum. The 100-ms window of maximum effect size was defined as this latency plus and minus 50 ms.

For Figures 2 to 6, we averaged the potential in the 100-ms time window selected above. For speed of processing and to align with standard ERP analyses practices, epochs of 1 second (from −0.3 to 0.7 seconds) instead of 3 seconds were extracted (except for section A.5). Since no baseline period is removed, the length of the data epochs does not influence the data quality measures (except when continuous data portions are rejected, in which case more 1-second epochs than 3-second epochs may be extracted). For each channel and each subject, we drew 50 randomly selected epochs (with replacement) from each condition and calculated the percentage of significant channels between conditions (p<0.05) using an unpaired t-test (*ttest_cell* function of EEGLAB). We repeated this procedure 20,000 times to obtain the average percentage of significant channels for each subject (one value per subject).

When comparing two preprocessing methods, our null hypothesis was that there was no difference between the two methods. We calculated the pairwise difference between methods (using subjects as cases) and computed confidence intervals and significance using 20,000 bootstraps (this is equivalent to using a paired t-test, although, unlike a t-test, it does not require the data to be normally distributed). When a rejection method removed too many trials for some subjects (see below), we ignored them when calculating significance.

Some of the methods involved data cleaning, which flagged bad data trials. For these methods, since significance is affected by the number of trials, we bootstrapped the number of remaining data trials (instead of 50) to compute the percentage of significant channels - except in the Discussion where we bootstrapped 50 trials to assess whether rejection methods were able to extract a set of good trials (and most were). When using all remaining data trials to compute significance, we checked for saturation of our performance metric, and that not all electrodes were significant. For both the Go/No-go and the Oddball datasets, about half of the subjects reached 80% significant channels with high-passed filtered data. Still, none reached 100%, and even when this value was above 90% (only four such subjects in all datasets), the EEGLAB pipeline could sometimes improve performance. If the percentage of bad data trials for a given condition and a given subject was more than 75% of the total number of trials, we ignored the subject. When comparing methods, significance was not calculated when the number of remaining subjects was less than 4.

We rounded all significant effects to the nearest percent. P-values are reported uncorrected, but we do not report any significant p-value above 0.01 – p-values between 0.01 and 0.05 are reported as trends - accounting for Bonferroni type correction for multiple comparisons when the number of comparisons in a plot is less than 5.

#### Computing platform

We performed all computations on Expanse, a high-performance computing resource at the San Diego Supercomputer Center part of the Neuroscience Gateway.^27^ On Expanse, we used MATLAB 2020b and Python 3.8. Except when troubleshooting, all computations were run in non-interactive mode using the SLURM workload manager. We used a total of approximately 10,000 core hours to run our comparisons.

### Data

#### Data availability

All data is publicly available on the OpenNeuro.org and NEMAR.org web platform under the following public digital object identifiers doi:10.18112/openneuro.ds002680.v1.2.0, doi:10.18112/openneuro.ds002718.v1.0.5, doi:10.18112/openneuro.ds003061.v1.1.1.

#### Go/No-go dataset

This experiment is a go-no visual categorization task where 14 participants responded when they saw an animal in a briefly flashed photograph. Fifty percent of the photographs contained images of animals photographed from different angles, and the other 50% did not contain images of animals or humans. Images were presented animal for 20 ms at random intervals of 1.8–2.2 s. We compared correct targets - go responses on animals - to correct distractors - no-go responses on non-animal images.^28^ To reduce the amount of data to process, we only used data from one of the two recording sessions (session 1), which amounted to an average of 228 targets (standard deviation: 17 targets) and 235 distractors (standard deviation: 10 distractors). The data was acquired at 1000 Hz and resampled to 250 Hz for subsequent analysis using the MATLAB *resample* function.

#### Face dataset

This multi-modal dataset^11^ contained EEG data acquired by an Elekta MEG machine and has been used in many publications, including the comparison of open-source software packages.^29^ Eighteen participants judged the symmetry of facial photographs presented sequentially. A fixation cross was presented for a random duration between 400 and 600 ms, after which the stimulus (face or scrambled face) was superimposed for a random duration between 800 and 1,000 ms. During the 1,700 ms interstimulus interval a central white circle was shown. This task was accessory and was used to maintain participant engagement, as the goal of the experiment was to compare different types of faces: familiar, unfamiliar, and scrambled. Here we compared familiar and scrambled faces. We used a release of this dataset tailored for EEG research, where the six runs for each subject were concatenated, and the data was resampled at 250 Hz.^30^ Each subject saw 150 familiar and 150 scrambled faces. This dataset contained four non-EEG channels, which we ignored in all analyses.

#### Oddball dataset

In this experiment, 13 participants responded by pressing a button to infrequent oddball sounds (70 ms of 1000 Hz; 15% of the trials) interspaced by frequent standard sounds (70 ms at 500 Hz; 70% of the trials) and infrequent white-noise sounds (70 ms duration; 15% of the trials). Inter-stimulus interval was selected randomly from 0.9 to 1.1 seconds. Of the three data blocks (or runs) for each subject, only one run is analyzed here to speed up computation. Run 1 is truncated for one of the subjects, so we processed run 2 instead. We only considered correct responses to oddball targets and standard non-targets (behavioral response on oddball stimuli and absence of response on standard stimuli). There was an average of 92 correct targets (standard deviation: 25) and 518 correct non-targets (standard deviation: 8). The data were digitally resampled to 256 Hz before being processed using the BIOSEMI resampling program. This dataset contained 15 auxiliary channels, which were ignored in all analyses, except for the referencing (five of these auxiliary channels are used to re-reference the data in Figure 3: left and right mastoids, left and right earlobes, and nose).

### Filtering

As a reference filter in Figure 2, we used the *pop_basicfilter* method of ERPLAB v9.0,^31^ which allows designing a stable filter down to 0.01 Hz (Butterworth of order 4 with default ERPLAB roll-off parameter values). For Supplementary Figure 2, for EEGLAB, we used the default Hamming windowed sinc FIR filter of the Firfilt plugin (v2.6) with 0.5 Hz high-frequency cutoff and default parameters (the order of the filter was automatically calculated based on the signal sampling frequency). This filter was also used for all analyses of sections A.2-6. For MNE, we used the default filter (linear FIR filter of the *filter* method of the *raw* object with a high-pass filter at 0.5 and no low-pass filter – all other parameters set to defaults). For Brainstorm, we used the *process_bandpass* function with a FIR high-pass filter at 0.5, no low pass, and all default parameters (2019 FIR filter version, which is the default). For FieldTrip, we used the default settings for high-pass filtering (4th order Butterworth filter). For FieldTrip, we use the one-step procedure in the *ft_preprocessing* function, which filters data epochs instead of filtering the raw data and then extracting data epochs (“FieldTrip But epochs” in Supplementary Figure 2). Because of poor performance for the Auditory Oddball dataset, and upon contacting the FieldTrip tool developers, we also used a multi-step approach to first filter the data and then extract epochs using the *ft_redefinetrial* function (“FieldTrip But” in Supplementary Figure 2). All filters were non-causal two-pass filters (forward and backward); when present, the filter order indicated above refers to one of the two passes.

### Line noise rejection

For the notch linear filter, we used the *pop_eegfiltnew* function of EEGLAB 2022.1 with passband edges 48 and 52 and default parameters (FIR filter of order 415 for 250 Hz data, 2 Hz transition bandwidth, zero phase, non-causal, cutoff frequencies at - 6bD of 49 to 51 Hz). For the Notch filter, we used the IIRFilt EEGLAB plugin (v1.03) with 48 and 52 and default parameters (IIR filter of order 12 and transition bandwidth of 1 Hz). For rejecting channels containing line noise, we used the *lineNoiseCriterion* parameter (with all other parameters disabled) of the *clean_rawdata* plugin of EEGLAB (v2.7) with a range of standard deviation thresholds (see main text). We also tested the *cleanline* EEGLAB plugin (v2.0), which adaptively estimates and removes sinusoidal (e.g., line) noise from scalp channels using multi-tapering and a Thompson F-statistic, leveraging methods developed in the Chronux toolbox.^13^ We finally used the *Zapline-plus* (v1.2.1) EEGLAB plugin, a method that combines spectral and spatial filtering to remove line noise.^14^ We used default settings for both *cleanline* and *Zapline-plus* for the 50-Hz frequency band.

### Re-referencing

The auditory Oddball dataset was recorded with additional channels to test re-referencing methods: mastoids, earlobes, and nose. These additional channels are recorded in the auxiliary channels of the BIOSEMI amplifier (EXG 1-5) and are not included in any other analyses. EEGLAB (v2022.1) *pop_reref* method was used to reference the data to these channels. We also used the PREP pipeline *performReference* function (with default parameters), which calculates a robust reference using the RANSAC algorithm.^23^ We used the EEGLAB *pop_reref* function and the FieldTrip *ft_preproc_rereference* function to calculate the average reference with default parameters. We also used the FieldTrip *ft_preproc_rereference* function to compute the median reference. For the REST reference^15^ and following the FieldTrip tutorial (https://www.fieldtriptoolbox.org/example/rereference/; August 8, 2022), we used the same 10- to 20-template montage (Easycap-M1) for all datasets, and calculated a leadfield matrix for the 4-sphere spherical head model (conductances of 0.33, 1, 0.0042, and 0.3300; radii of 71, 72, 79, and 85 mm). Finally, still using FieldTrip, we calculated the longitudinal and circumferential daisy chain references, which extract a small subset of channels and calculate their pairwise difference.

### Automated artifact rejection

#### EEGLAB clean_rawdata channel rejection

The *clean_rawdata* EEGLAB plugin (v2.7) was used for detecting bad channels with correlation thresholds ranging from 0.15 to 0.975 (see Table 1). All other data rejection methods of *clean_rawdata* were disabled. Channels below the correlation threshold are interpolated using spherical splines (*pop_interp* function of EEGLAB 2022.1).

#### EEGLAB clean_rawdata ASR rejection

The *clean_rawdata* EEGLAB plugin (v2.7) was used for detecting bad segments of data with thresholds ranging from 5 to 200 (see Table 1). This plugin uses the Artifact Subspace Reconstruction method^15^ to detect and correct bad portions of data. We only used the artifact detection method of ASR to remove bad segments of data (which is the default in EEGLAB for offline processing) and did not correct them. All other data rejection methods of *clean_rawdata* were disabled.

#### EEGLAB ICLabel eye movement and muscle rejection

ICA was performed using Infomax (*Picard* plugin v1.0) and default parameters in EEGLAB 2022.1 (EEGLAB automatically sets the maximum iteration of *Picard* to 500 instead of 100). *ICLabel* (v1.4)^16^ is a machine learning algorithm able to detect artifactual ICA components based on their topography and activity. Each component is assigned a probability of belonging to 1 of 7 classes, which include the muscle and eye movement artifact classes. We applied the *ICLabel* default method to detect eye and muscle artifacts with probability thresholds ranging from 0.5 to 0.9. When testing other ICA algorithms (*runica, AMICA, FastICA, sobi*; Supplementary Figure 8), we use their implementation in EEGLAB or EEGLAB plugins with default parameter values.

#### FieldTrip ft_artifact_zvalue artifact parameters

We used the function *ft_artifact_zvalue* (FieldTrip version of August 7, 2022) to remove artifacts. We followed the online tutorial (https://www.fieldtriptoolbox.org/tutorial/automatic_artifact_rejection/; version of August 8, 2022) and used the code snippets to remove EOG and muscle artifacts. For detecting eye movements, we used frontal channels AF7, AF3, Fp1, Fp2, and Fpz (Oddball dataset), AF3, AF4, AF7, AF8, and Fpz (Face datasets), channels Fp1 and Fp2 (Go/No-go dataset). We left all other parameters as in the tutorial (*trlpadding*=0; *artpadding*=0.1; *fltpadding*=0; 4th order Butterworth filter and use of the Hilbert method). We used a frequency range of 2 to 15 Hz, as recommended in the FieldTrip tutorial, and varied the z-score threshold from 1 to 6. For detecting muscle artifacts, we changed the channels to T7, T8, TP7, TP8, P9, and P10 for the Oddball and Face datasets and T5, T6, CB1, and CB2 for the Go/No-go dataset. We used the default 9-order Butterworth filter and boxcar parameter to 0.2, but changed the frequency range from 100 to 110 Hz (the upper edge of this filter is lower than in the FieldTrip tutorial to accommodate our lower data sampling rate) and varied the z-score threshold from 1 to 6. Note that z-score calculated on a subset of channels are likely more relevant for MEG than for EEG: For EEG, the number of channels with noise might depend on the reference. High frequency noise is also likely stronger in MEG than EEG where it might be smeared out.

#### Brainstorm bad segment detection

We used the Brainstorm (version of August 5, 2022) functionality to detect bad portions of data (menu item “Detect other artifacts” corresponding to the command line function *process_evt_detect_badsegment*). We varied sensitivity from 1 to 5 (the only values allowed) for the low-frequency and high-frequency artifact detection. We also used the Brainstorm function to detect bad data trials (*process_detectbad*) and varied the threshold from 200 to 5000 (Table 1).

#### MNE Autoreject parameters

We used MNE 1.1.0 on Python 3.8 with default scientific and plotting libraries and MNE libraries *eeglabio* 0.0.2 and *Autoreject* 0.3.1. We could not find any function to automatically reject bad portions of data or bad channels within MNE itself. The *Autoreject*^32^ plugin allows the correction and rejection of bad data. Although it is not officially part of MNE, it was made by MNE core developers. Upon contacting other MNE core developers, no alternative method was proposed. Unlike other artifact rejection methods with tunable parameters, *Autoreject* automatically scans the parameter space to reject the best number of channels and bad data regions. The algorithm was validated on four EEG datasets^32^ and has been used with default parameters by at least one other group.^33^ We followed the online documentation (https://autoreject.github.io/stable/auto_examples/plot_autoreject_workflow.html; August 8^th^ 2022) and used, as in the tutorial, 1 to 4 channels to interpolate. Another parameter in the *Autoreject* tutorial is the number of epochs to fit. The proposed method uses the first 20 epochs to speed up computation. We tried using the first 20 epochs and all epochs (Table 1).

### Pipelines

#### Common processing for all pipelines

High-pass filtering at 0.5 Hz was applied using the default method in each software package (Finite Impulse Response for EEGLAB, MNE, and Brainstorm and Butterworth filter for FieldTrip; see also the Filtering Method section).

#### HP 0.5 Hz pipeline

We use the default EEGLAB FIR 0.5-Hz high-pass filter and no other preprocessing. See the Filtering Method section.

#### EEGLAB pipeline

We use the default FIR 0.5-Hz high-pass filter, followed by electrode line noise detection and interpolation (4 standard deviation threshold; section A.4), *clean_rawdata* channel correlation removal (0.9 correlation threshold), *clean_rawdata* ASR rejection (threshold of 20), ICA followed by *ICLabel* with 0.9 probability thresholds for muscle and eye.

#### FieldTrip pipeline

We used the default Butterworth 0.5 Hz high-pass filter. FieldTrip one step *ft_preprocessing* function filters data epochs instead of filtering the raw data then extracting data epochs, leading to poor performance (Supplementary Figure 2), so we used a multi-step approach to first filter the data then extract epochs using the *ft_redefinetrial* function. We rejected data epochs with high and low-frequency artifacts with a 4-dB threshold, which is the default in the FieldTrip tutorial (see the section *“FieldTrip ft_artifact_zvalue artifact parameters”* and section A.4).

#### Brainstorm pipeline

We used the default Brainstorm 0.5-Hz high-pass filter and rejected low- and high-frequency artifacts with sensitivity level 5 (section A.4). We rejected bad trials with a threshold of 200 (section A.4).

#### MNE pipeline

We used the default MNE 0.5-Hz high-pass filter and rejected and repaired artifacts using *Autoreject* (20 trials used for the fitting process; Section A.4).

#### HAPPE pipeline

HAPPE^34^ is an EEGLAB-based processing pipeline that uses the MARA EEGLAB plugin for rejecting artifacts using ICA^26^ and the FASTER EEGLAB plugin to interpolate bad data segments.^25^ It is an integrated pipeline, although it allows users to set a peak-to-peak raw data threshold in microvolts. We tried a range of 50 to 150 microvolts for this threshold in 10 increments but did not find a significant difference between thresholds or with no threshold, so we set the threshold to 100. We used the default setting for rereferencing, which computes an average reference (not re-referencing is not an option). We modified the core code to decrease the high-pass filter cutoff frequency to 0.5 Hz (instead of the 1-Hz default) to match the first preprocessing step with other pipelines. We disabled the *cleanline* plugin to remove line noise because it produced errors on some datasets. We made other minor modifications to be able to process the Go/No-go, Face, and Oddball datasets and issued version 2.0 of the HAPPE pipeline for others to use (https://github.com/arnodelorme/happe). Unlike other pipelines, this pipeline automatically re-reference the data which may explain its poor performance in Figure 5.

## Acknowledgment

This work was supported by NIH (R01NS047293-18 and R24MH120037-04). The author wishes to thank Robert Oostenveld and Johanna Wagner for their comments, Amitava Majumdar and his team for time allocation at the San Diego Supercomputer Center, and Anne Stanford for proofreading the manuscript.

