## Supplementary material for "EEG is better left alone"

| | Visual Go/No-go EEG dataset | Face EEG dataset | Auditory oddball EEG dataset |
|---|---|---|---|
| **Public DOI** | doi:10.18112/openneuro .ds002680.v1.2.0 | doi:10.18112/openneur o.ds002718.v1.0.5 | doi:10.18112/openneuro.ds0 03061.v1.1.1 |
| **Acquisition hardware** | Neuroscan Synamps 5083 | Elekta Neuromag Vectorview 306 | ActiveTwo BIOSEMI |
| **Hardware reference** | Cz | Noze | Internal (CMS/DRL placed on each side of POz) |
| **Number of subjects** | 14 | 18 | 13 |
| **Sensory modality** | Visual | Visual | Auditory |
| **Number of trials per conditions** | 228±17 targets<br>235±10 distractors | 150 familiar faces<br>150 scrambled faces | 92±25 oddball<br>518±8 standard |
| **Inter-stimulus interval** | 1.8 to 2.2 seconds | 2.9 to 3.3 seconds | 0.9 to 1.1 seconds |
| **Power line frequency** | 50 Hz (notched)* | 50 Hz | 50 Hz |
| **Hardware high-pass** | DC† | 0.1 Hz† | 0.016 Hz† |
| **Number of EEG channels** | 31 | 70 | 64 |
| **Montage type** | 10-20 & custom | 10-20 digitized | 10-20 |
| **Original sampling rate** | 1000 | 1100 | 2048 |
| **Sampling rate for analysis** | 250 | 250 | 256 |

*The notch filter is likely a digital filter applied on the Synamps hardware. It might be applied to data acquired at a higher sampling frequency than the acquisition frequency. This information was not available in the 2007 SynAmps manual.
†The 2007 SynAmps manual mentions that the Neuroscan amplifier is a DC amplifier, and the analog high-pass cutoff frequency is not indicated. The Elekta 2005 manual page 42 mentions a default value for EEG high-pass at 0.1 Hz. The BIOSEMI amplifier applied a low 3dB attenuation analog high-pass filter (0.016 Hz to 0.16 Hz are the edge bands), and DC may be recovered.

*Supplementary Table 1. Description of three experiments used to assess data quality: Go/No-go (visual categorization), Face characteristics, and auditory oddball experiments. For each experiment, we compare two conditions. The first experiment (Go/No-go) is a go-no visual categorization task where participants respond when they see an animal in a briefly flashed photograph, and we compared correct animal targets (50%) to correct distractors (50%). In the second experiment, we compared familiar and scrambled faces presented to participants who performed an accessory task designed to maintain their attention. In the third experiment, participants had to respond to oddball sounds (70 ms of 1000 Hz) interspace by frequency standard sounds (70 ms at 500 Hz). The number of trials differed for each subject in the Go/No-go and Oddball task since we only considered correct responses (mean and standard deviation are indicated). The data is digitally resampled before being processed.*

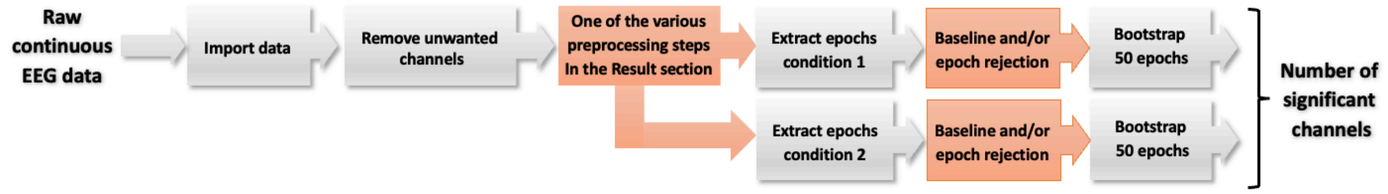


*Supplementary Figure 1. Schematic of data processing. For each dataset, the data is imported, then unwanted non-EEG channels are removed. Then a preprocessing step is applied (filtering, referencing, artifact rejection). The preprocessing step varies, and different methods are tested in this manuscript. Then data epochs are extracted for two conditions – the conditions depend on the dataset (see Supplementary Table 1). The pre-stimulus baseline activity may optionally be removed for each data epoch, and some epochs may be removed. 50 data epochs are then bootstrapped, and the number of significant channels is calculated. The last bootstrap step is repeated to obtain a confidence interval for the number of significant channels. Grey boxes indicate fixed processing steps, and orange boxes indicate processing steps that vary depending on the sub-section in the Result section.*

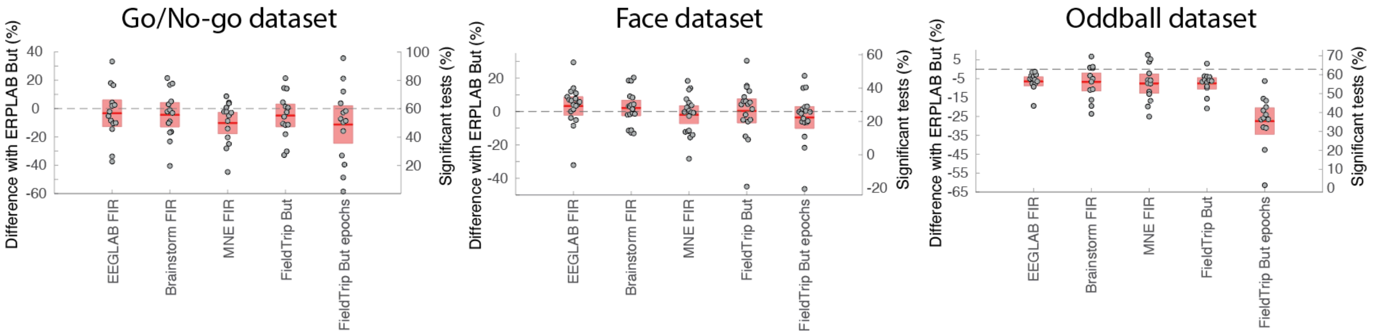


*Supplementary Figure 2. Influence of software implementation for 0.5 Hz high-pass filter (see Figure 2 for details regarding the ordinate scales, and the Methods section for filter implementation).*

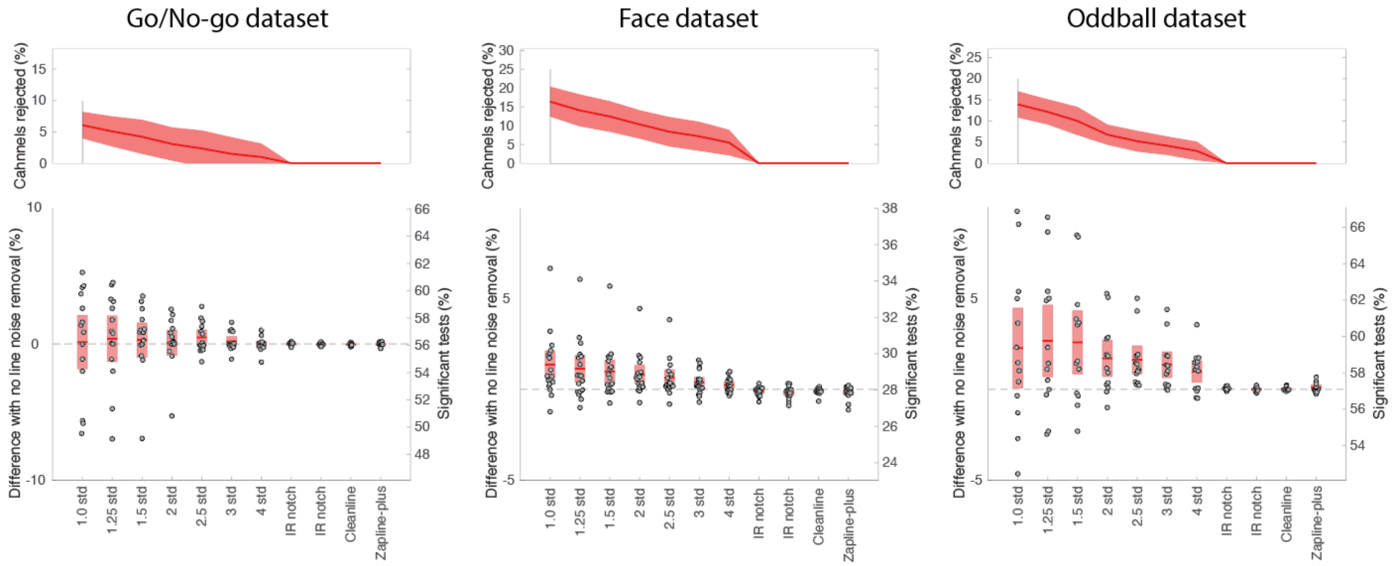


*Supplementary Figure 3. Influence of line noise removal methods on the number of significant channels compared to no filtering for three datasets. The top row shows the percentage of channels rejected. The bottom panels show the 95% confidence interval (red region) of the significant difference, and the dots indicate individual subjects (see Figure 2 for details regarding the ordinate scales).*

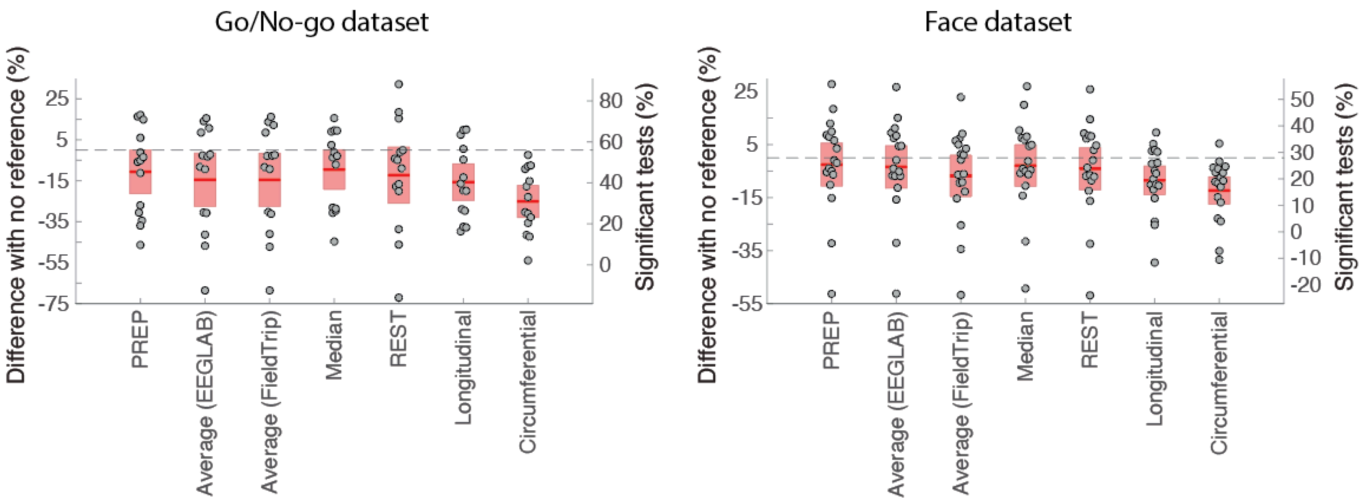


*Supplementary Figure 4. Influence of the reference on the number of significant channels compared to no reference for the Go/No-go (Cz hardware reference) and Face (nose hardware reference) datasets. See Figure 2 for details regarding the ordinate scales.*

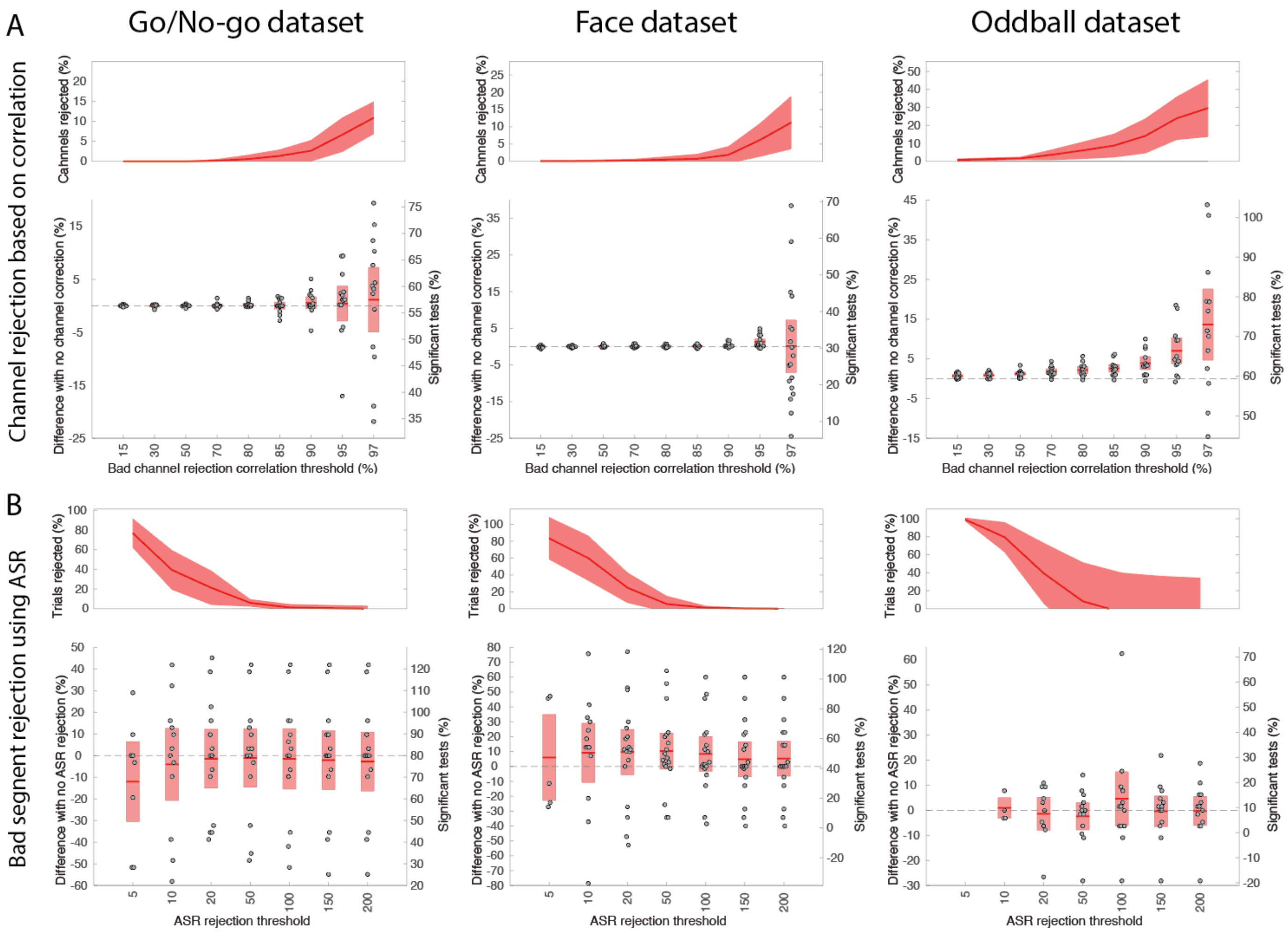


*Supplementary Figure 5. A. Clean_rawdata channel correlation rejection percentage of channel significant as a function of the correlation threshold for three datasets. B. Clean_rawdata channel ASR bad segment rejection as a function of the ASR threshold. Bottom box plots for each condition show the difference in the percentage of channels being significant for different z-score thresholds. The upper plots show the percentage of rejected channels and the standard deviation (shaded area). See Figure 2 for details regarding the ordinate scales.*

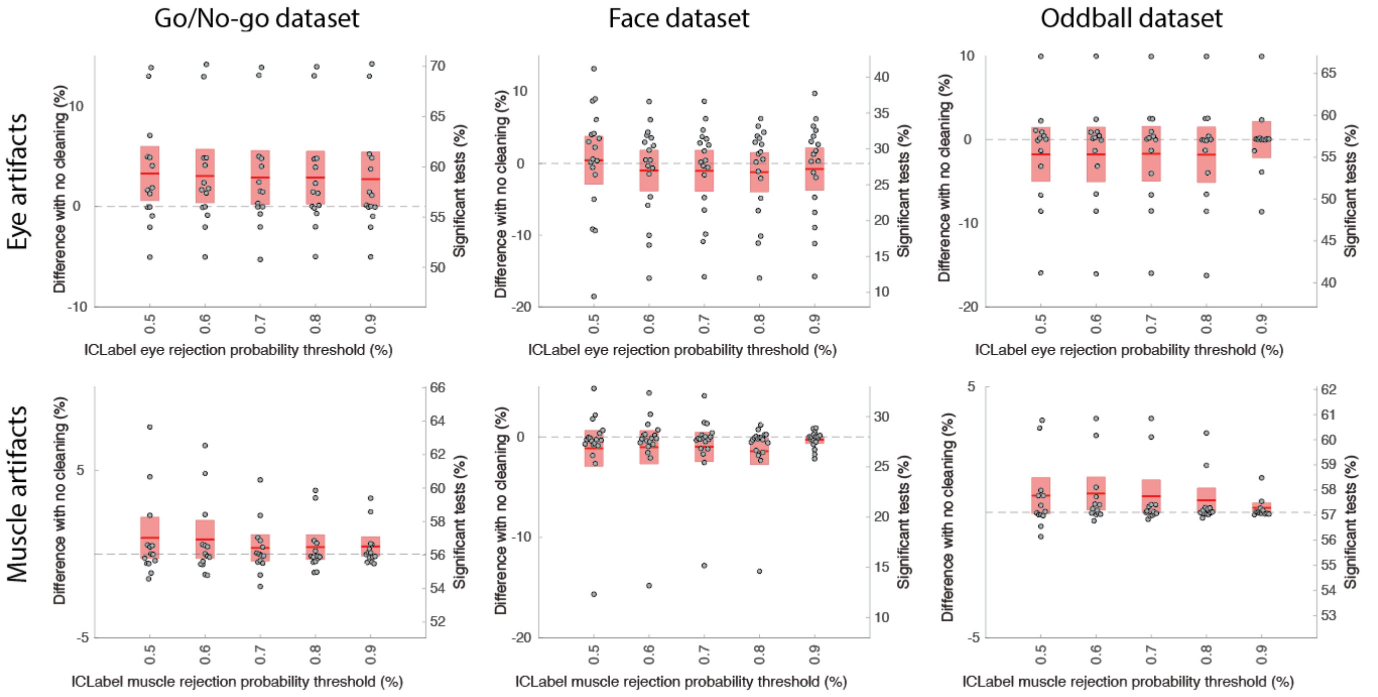


*Supplementary Figure 6. Performance of the ICLabel eye (top row) and muscle (bottom row) ICA component rejection for different likelihood thresholds on three datasets. ICLabel assigns a probability for each component to belong to the Eye or Muscle class. For a given threshold, all components with a likelihood of being Eye or Muscle artifacts above that threshold are rejected. See Figure 2 for details regarding the ordinate scales.*

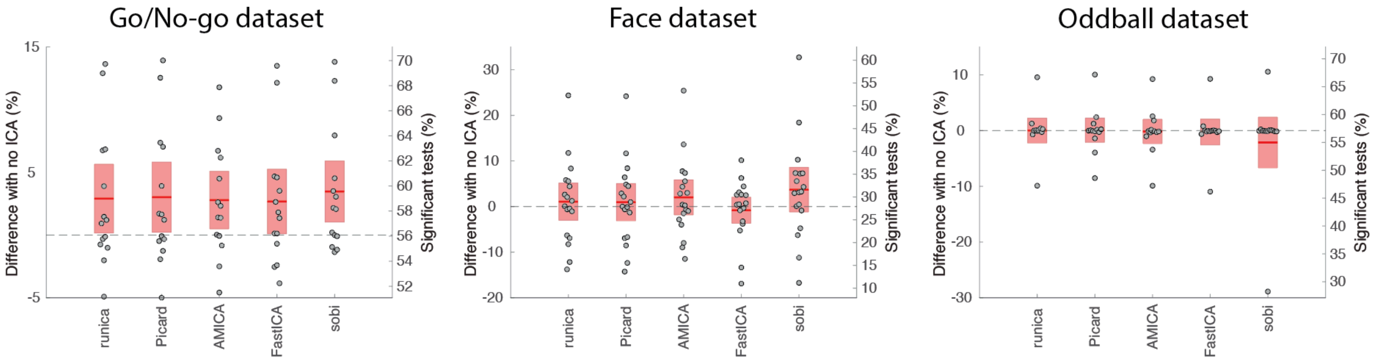


*Supplementary Figure 7. ICLabel applied to different ICA decomposition for three datasets compared to no rejection (see Figure 2 for details). Both eye and muscle components were rejected (threshold of 0.9; see Methods). See Figure 2 for details regarding the ordinate scales.*

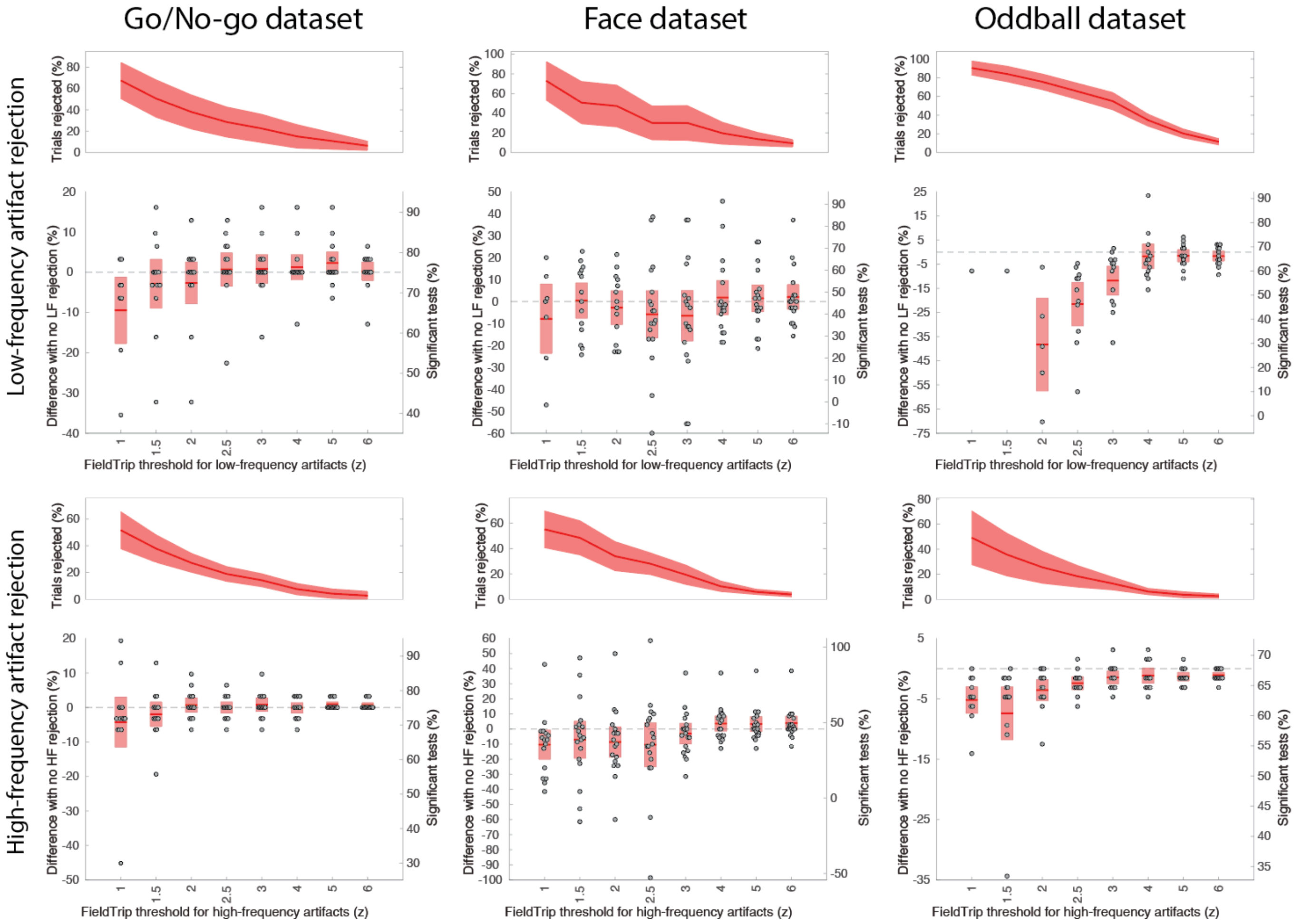


*Supplementary Figure 8. FieldTrip automated artifact rejection for low-frequency (top row) and high-frequency (bottom row) artifacts and for three datasets compared to no rejection on data high-pass filtered at 0.5 Hz. Bottom box plots for each condition show the difference in the percentage of channels being significant for different z-score thresholds (see Figure 2 for details regarding the ordinate scales). The upper plots show the percentage of rejected trials and the standard deviation.*

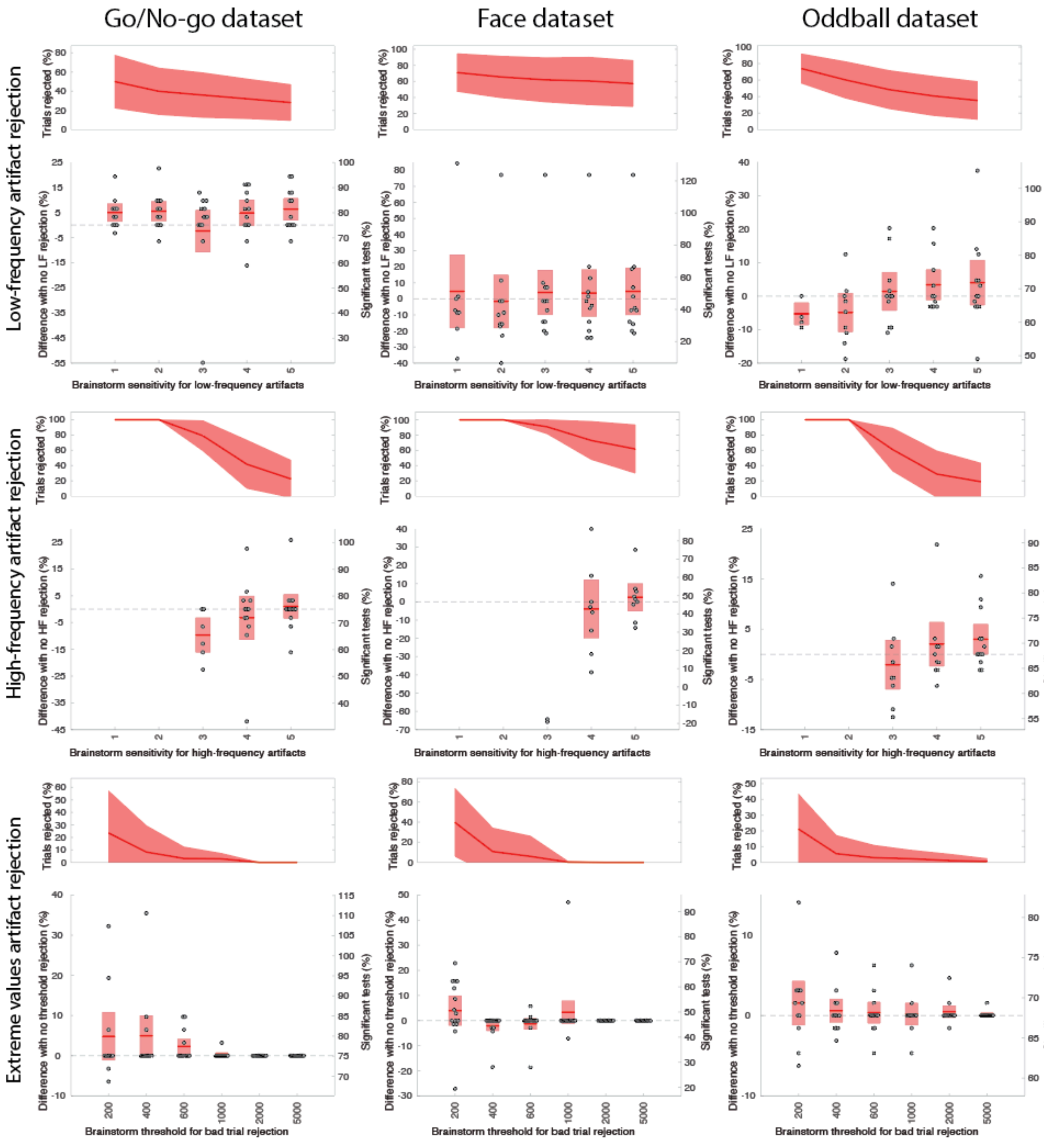


*Supplementary Figure 9. Brainstorm artifact rejection for low-frequency (top row), high-frequency (middle row) artifacts, and trials exceeding a threshold (bottom row) for three datasets compared to no rejection on data high-pass filtered at 0.5 Hz. The upper plots in each condition show the percentage of trials rejected, and the lower plots show the difference in the percentage of significant channels for different sensitivity levels*

*(low-frequency and high-frequency artifact rejection) and different thresholds for the extreme values artifact rejection method. See Figure 2 for details regarding the ordinate scales.*

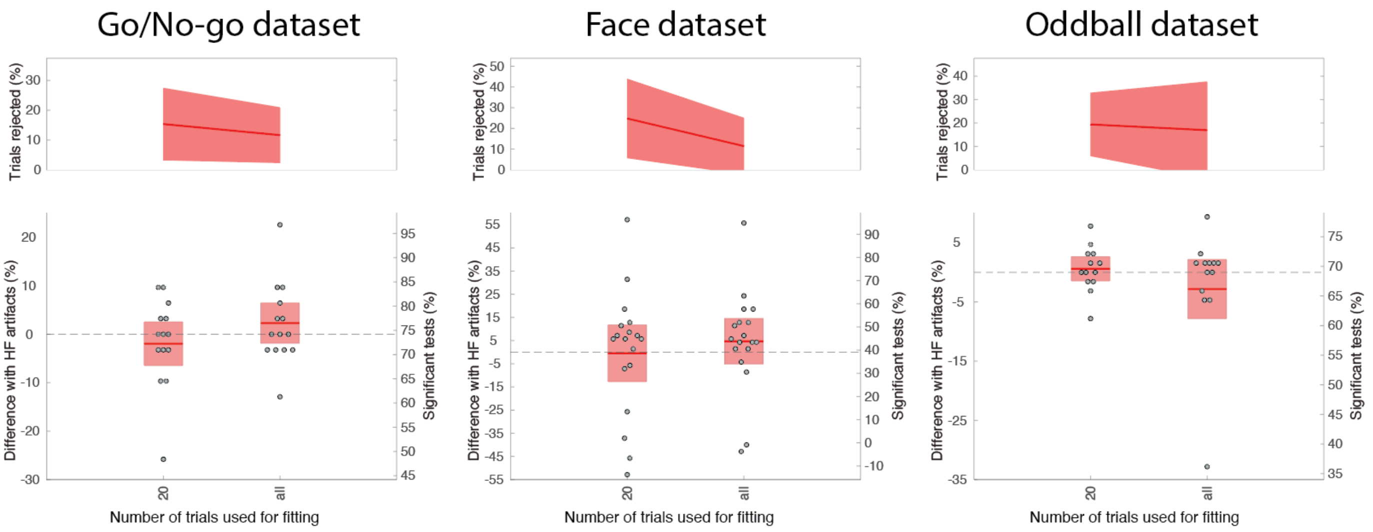


*Supplementary Figure 10. MNE Autoreject automatic for three datasets compared to no rejection on data high-pass filtered at 0.5 Hz. The upper plots in each condition show the percentage of trials rejected, and the lower plots show the difference in the percentage of significant channels when 20 or all trials are used for fitting the Autoreject model. See Figure 2 for details regarding the ordinate scales.*

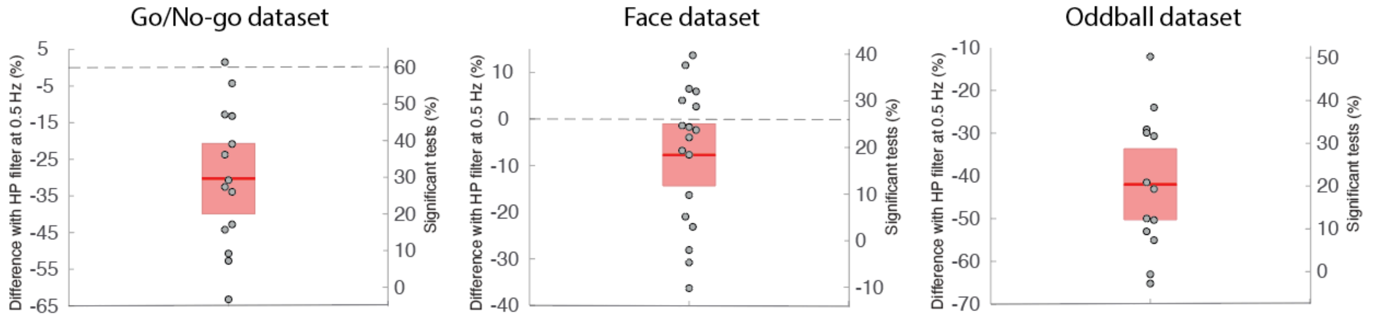


*Supplementary Figure 11. Difference in percentage of significant channels between using a filter at 0.01 Hz with a 200-ms pre-baseline interval and a filter at 0.5 Hz with no baseline (ordinate 0) for three datasets. See Figure 2 for details regarding the ordinate scales.*

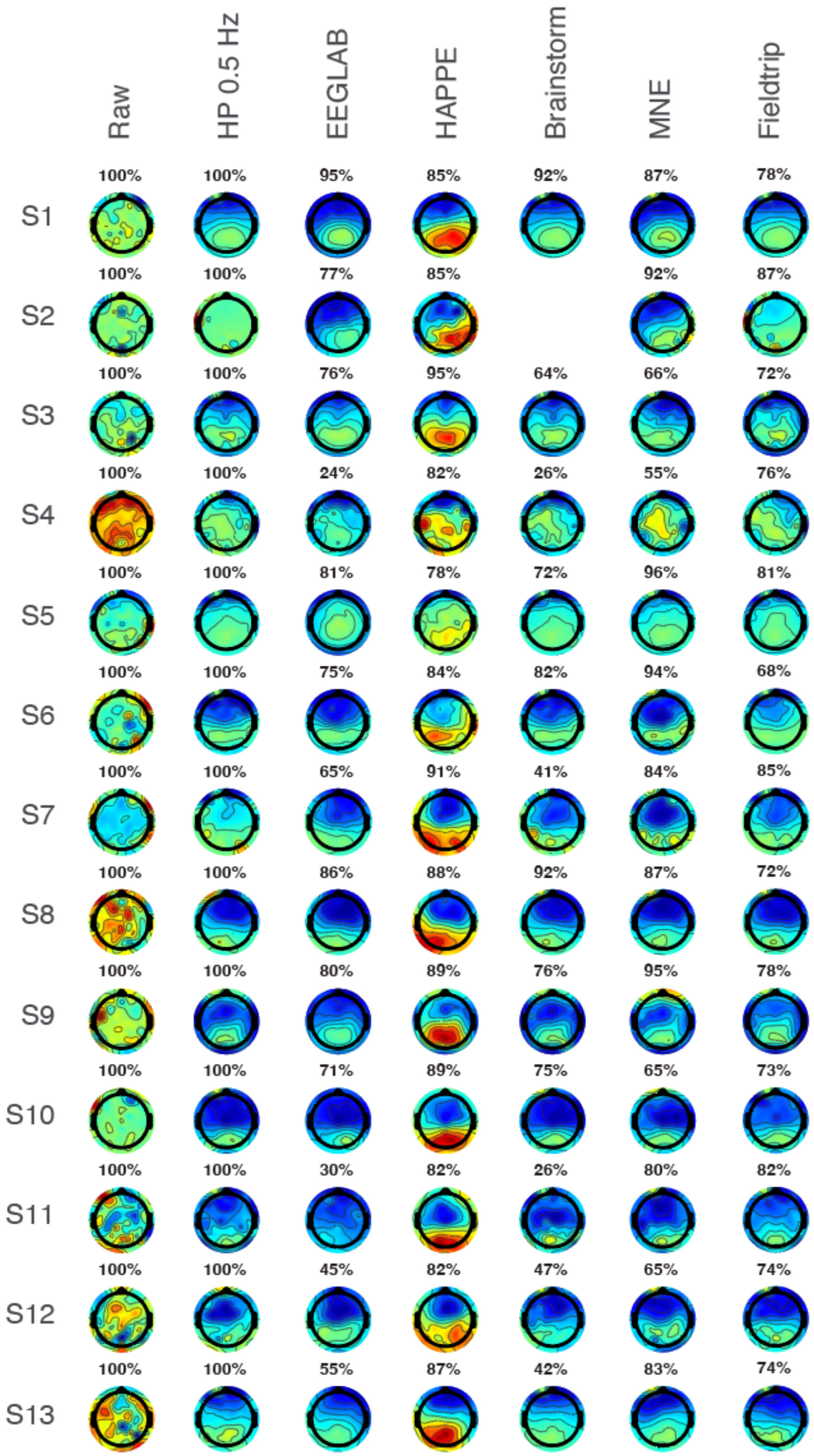

Raw
HP 0.5 Hz
EEGLAB
HAPPE
Brainstorm
MNE
Fieldtrip
S1
100% 100% 95% 85% 92% 87% 78%
S2
100% 100% 77% 85% 92% 87%
S3
100% 100% 76% 95% 64% 66% 72%
S4
100% 100% 24% 82% 26% 55% 76%
S5
100% 100% 81% 78% 72% 96% 81%
S6
100% 100% 75% 84% 82% 94% 68%
S7
100% 100% 65% 91% 41% 84% 85%
S8
100% 100% 86% 88% 92% 87% 72%
S9
100% 100% 80% 89% 76% 95% 78%
S10
100% 100% 71% 89% 75% 65% 73%
S11
100% 100% 30% 82% 26% 80% 82%
S12
100% 100% 45% 82% 47% 65% 74%
S13
100% 100% 55% 87% 42% 83% 74%

*Supplementary Figure 12. Scalp topography at 400-500 ms for the 13 subjects of the Oddball datasets and for different preprocessing pipelines. The number above each scalp topography indicates the percentage of remaining trials. Each scalp map is scaled to the minimum and maximum signal amplitude. Both the EEGLAB and HAPPE pipelines remove eye artifacts using ICA, which improves the aspect of scalp topography in most subjects over frontal regions. Brainstorm is missing a scalp topography for subject S2 because it removed all the data trials for one subject.*
